# Synergistic effects of Cyp51 isozyme-specific azole antifungal agents on fungi with multiple cyp51 isozyme genes

**DOI:** 10.1101/2025.04.16.649168

**Authors:** Masaki Ishii, Kazuki Ishikawa, Koji Ichinose, Takashi Yaguchi, Tsuyoshi Yamada, Shinya Ohata

## Abstract

Pathogenic fungi pose significant societal challenges. The limited availability of therapeutic targets due to the eukaryotic nature of fungi emphasizes the importance of available drug targets such as Cyp51, a crucial enzyme in ergosterol biosynthesis, inhibited by azole antifungals. This study explored the susceptibility patterns of azole antifungals against Cyp51 isozyme deletion strains (Δ*cyp51A* and Δ*cyp51B*) in *Trichophyton rubrum*, the predominant dermatophyte species. Distinct susceptible patterns were observed among azole antifungals for Δ*cyp51A* and Δ*cyp51B*. Although most azole antifungal agents exhibited increased antifungal activity against Δ*cyp51A*, select agents demonstrated increased antifungal activity against Δ*cyp51B*. Remarkably, fluconazole, sulconazole, and imazalil exhibited relatively increased activity against Δ*cyp51A*, whereas prochloraz demonstrated increased activity against Δ*cyp51B*. Combining these isozyme-selective agents exerted synergistic effects against the wild-type strain and the parent *ku80*-knockout strain but not against individual Cyp51 knockout mutants. Hence, the two Cyp51 isozymes, Cyp51A and Cyp51B, may be inhibited by distinct azole antifungals, exerting a synergistic effect with the dual azole antifungal combination. This synergistic effect was also observed on another fungal species, *Aspergillus welwitschiae*, which also has two Cyp51 isozymes. These data demonstrate that combining azole antifungals with different Cyp51 isozyme selectivities exerts synergistic effects against fungi possessing multiple Cyp51 isozymes. This study proposes a novel therapeutic approach for addressing fungal infections through the combination of antifungal drugs that inhibit the same enzymatic activity but exhibit different isozyme selectivity. It also emphasizes the potential for developing drugs targeting specific isozymes, a previously underutilized approach in the realm of antifungal drug development.

## INTRODUCTION

Fungal infections represent a significant global health challenge, causing 3.752 million annual deaths (1). Among nonlethal fungal infections, dermatophytosis affects >10% of the world’s population and significantly influences patients’ quality of life through direct symptoms such as itching and inflammation, and also potentially exacerbating conditions such as asthma and increasing the risk of developing diabetic foot ulcers (2–4). The development of selective antifungal drugs is particularly challenging due to the similarity between fungal and mammalian cellular components (5–7), necessitating the discovery of both novel drug targets and novel approaches to existing molecular targets.

Combination therapy typically uses drugs targeting distinct molecules to exert additive or synergistic effects (8). Nevertheless, studies have demonstrated that combining drugs from the same class can also be effective, as demonstrated by the synergistic effects of β-lactam antibiotic combinations both *in vitro* and in clinical settings (9–11). One explanation for the synergistic effects is that β-lactam antibiotics interact with multiple penicillin-binding proteins (PBPs) with varying affinities (12). When two β-lactams with different PBP preferences are combined, they can effectively inhibit multiple PBPs at lower concentrations. Although these synergistic effects have been well investigated in antibacterial therapy (10, 11, 13), there is a lack of similar investigations with antifungal drugs.

Azole antifungals, the most widely used class of antifungal drugs, target the sterol 14α-demethylase enzyme Cyp51 in the ergosterol biosynthesis pathway (14). Although Cyp51 is encoded by a single essential gene (*erg11*) in *Candida albicans* and *Saccharomyces cerevisiae*, some filamentous fungi, including *Trichophyton* and *Aspergillus* species, possess multiple Cyp51 genes (14). For instance, *Trichophyton rubrum*, the most common causative agent of dermatophytosis, expresses both Cyp51A and Cyp51B (14). Nonetheless, the relative specificity of existing azole antifungal agents for these isozymes remains unexplored.

We hypothesized that different azole antifungals exhibit varying sensitivities for these isozyme deletion mutants and that combinations of azoles with distinct preferences exert synergistic effects. In this study, we generated *cyp51A* and *cyp51B* deletion strains in *T. rubrum* to explore the isozyme selectivity of various azole antifungals. Our results showed that different azoles in fact exhibit selective activity against each Cyp51 isozyme. We also demonstrated synergistic antifungal activity using the combination of imazalil and prochloraz, which exhibited the highest increases in susceptibility for Δ*cyp51A* and Δ*cyp51B* strains, respectively. These findings suggest a novel strategic approach to improve antifungal therapy.

## RESULTS

### Characterization of Cyp51 Isozyme Deletion Strains in T. rubrum

We recently generated a strain deficient in *ku80*, which is involved in nonhomologous end joining, for efficient genetic recombination in *T. rubrum* (15). Based on that report, we generated deletion strains of individual Cyp51 isozymes (Cyp51A and Cyp51B) in *T. rubrum* Δ*ku80* strain. The successful deletion of *cyp51A* and *cyp51B* was confirmed by Polymerase Chain Reaction (PCR) using the primers depicted in Fig. 1A and B (Fig. 1A–D). The *cyp51A* deletion strain (Δ*cyp51A*) demonstrated growth comparable to that of the parent strain Δ*ku80*, consistent with our previous observations of mutants (15). In the earlier study, the insertion of a drug resistance gene into the 3′-UTR region of *cyp51A* resulted in reduced expression without exhibiting any visible growth defects in *T. rubrum* (15). Interestingly, the *cyp51B* deletion strain (Δ*cyp51B*) exhibited a significant mycelial growth defect (Fig. 1E and F), which was consistently observed in multiple independently generated deletion strains (data not shown). This finding suggests that Cyp51B is essential for normal mycelial growth in *T. rubrum* under standard culture conditions.

**Figure 1.**
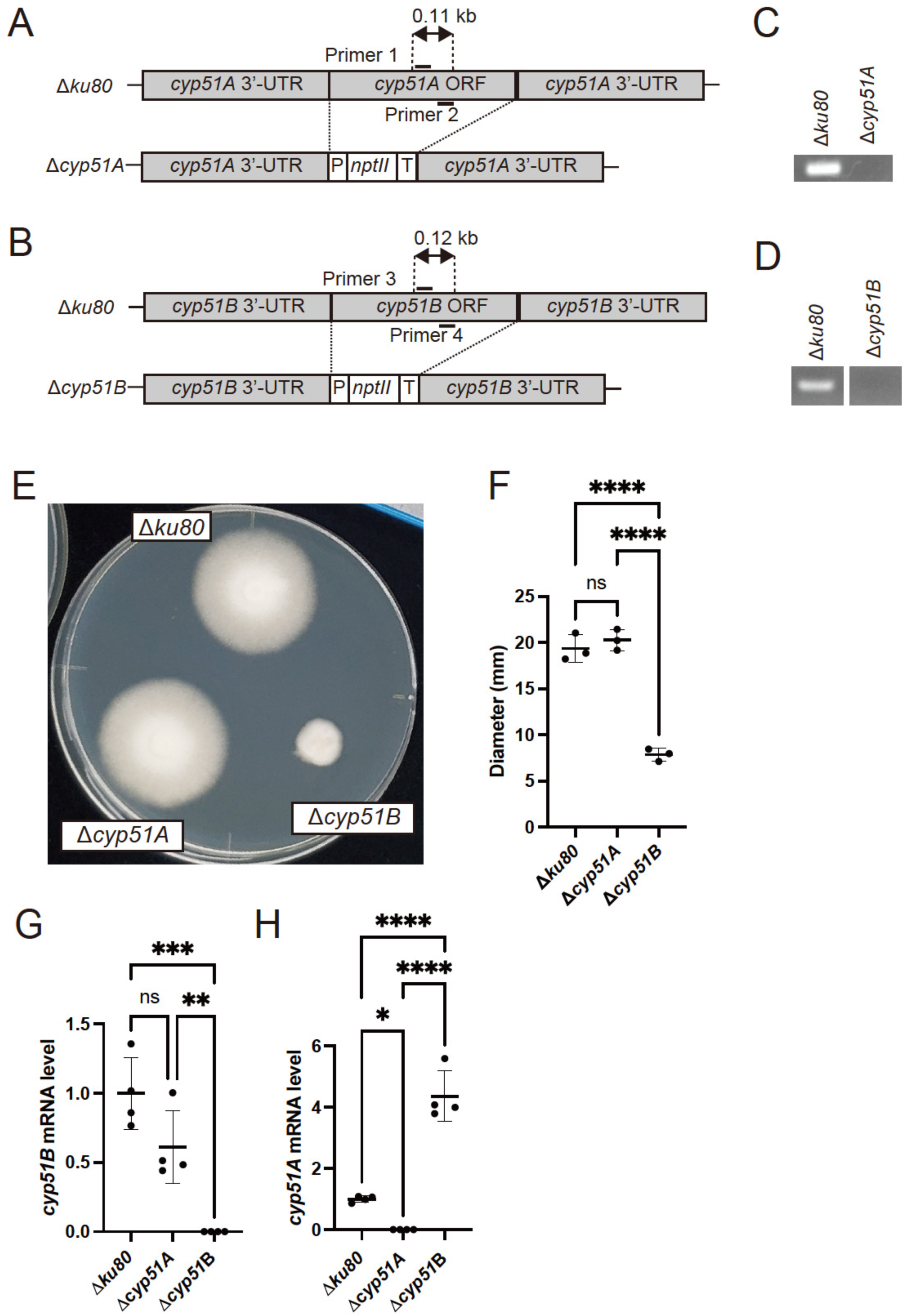
Generation and characterization of *cyp51A* and *cyp51B* deletion strains of *T. rubrum*. (A) Schematic of the *cyp51A* locus of WT and Δ*cyp51A* strains. (B) PCR analysis of total DNA samples from the Δ*cyp51A* strain. The fragments were amplified using primer pairs (Primer 1 and 2). Δ*ku80* was used as a control. (C) Schematic of the *cyp51B* locus of WT and Δ*cyp51B* strains. (B) PCR analysis of total DNA samples from the Δ*cyp51B* strain. The fragments were amplified using primer pairs (Primer 3 and 4). Δ*ku80* was used as a control. (E) Mycelial growth of Δ*ku80*, Δ*cyp51A*, and Δ*cyp51B* on SDA at 28°C for 14 days. (F) Colony diameter of Δ*ku80*, Δ*cyp51A*, and Δ*cyp51B* strains on SDA at 28°C for 14 days. The bars represent the standard deviation of data obtained from three independent experiments. (G-H) Expression levels of *cyp51B* (G) and *cyp51A* (H) mRNA in Δ*ku80*, Δ*cyp51A,* and Δ*cyp51B* strains as determined by qRT-PCR. The fold change represents the gene expression level compared with that of Δ*ku80*. The bars represent the standard deviation of data obtained from three independent experiments. \**P* < 0.05; \*\**P* < 0.01; \*\*\**P* < 0.001; \*\*\*\**P* < 0.0001.

Expression analysis revealed no compensatory upregulation of *cyp51B* mRNA when *cyp51A* was deleted, indicating that basal *cyp51B* mRNA expression is sufficient for normal growth in nutrient media (Fig. 1G). Nevertheless, *cyp51A* mRNA expression was upregulated in the *cyp51B* deletion strain (Fig. 1H). Moreover, exposure to azoles induced *cyp51A* mRNA expression but not *cyp51B* mRNA expression (15), suggesting that Cyp51A functions as an inducible isozyme that responds to insufficient ergosterol synthesis.

### Differential Sensitivity of Cyp51 Isozyme Deletion Mutants to Azole Antifungals

To gain insights into the Cyp51 isozyme selectivity of antifungal agents, we determined the MIC for 17 clinical antifungals and 13 pesticides against Δ*cyp51A* and Δ*cyp51B* strains. The Δ*cyp51A* strain demonstrated 2– to 256-fold decreases in MICs for most azole antifungals (27/30), except for posaconazole, lanoconazole, and luliconazole (Table 1). The increased drug susceptibility in the Δ*cyp51A* strain harboring only the Cyp51B isozyme suggests that most azole antifungal drugs are more selective against Cyp51B than against Cyp51A. The Δ*cyp51B* strain exhibited a more than 4-fold reduction of MICs only for isavuconazole and prochloraz but exhibited increased MICs for isoconazole, miconazole, neticonazole, oxiconazole, and sulconazole (Table 1). These data indicate that Cyp51A primarily mediates the natural tolerance to most azole antifungals in *T. rubrum* and that different azoles may exhibit distinct selectivity for Cyp51 isozymes.

**Table 1.** MIC values (μg/ml) and MIC ratio (MIC of Δ*ku80*/MIC of Δ*cyp51A* or Δ*cyp51B*) of antifungal drugs and pesticides against Δ*ku80*, Δ*cyp51A*, and Δ*cyp51B* strains.

| | MIC<sub>100</sub> ($\mu\text{g/ml}$) | | | MIC of $\Delta ku80$ / | MIC of $\Delta ku80$ / |
| --- | --- | --- | --- | --- | --- |
| | $\Delta ku80$ | $\Delta cyp51A$ | $\Delta cyp51B$ | MIC of $\Delta cyp51A$ | MIC of $\Delta cyp51B$ |
| Triazole |  |  |  |  |  |
| Efinaconazole | 0.02 | 0.005 | 0.01 | 4 | 2 |
| Itraconazole | 1 | 0.5 | 0.5 | 2 | 2 |
| Ravuconazole | 0.08 | 0.04 | 0.04 | 2 | 2 |
| Fluconazole | 32 | 0.5 | 16 | 64 | 2 |
| Voriconazole | 0.031 | 0.0078 | 0.016 | 4 | 2 |
| Posaconazole | 0.5 | 0.5 | 0.5 | 1 | 1 |
| Isavuconazole | 0.063 | 0.031 | 0.016 | 2 | 4 |
| Imidazole |  |  |  |  |  |
| Bifonazole | 0.52 | 0.065 | 1 | 8 | 0.5 |
| Clotrimazole | 0.26 | 0.13 | 0.13 | 2 | 2 |
| Isoconazole | 0.26 | 0.13 | 0.52 | 2 | 0.5 |
| Ketoconazole | 1.4 | 0.18 | 1.4 | 8 | 1 |
| Laniconazole | 0.00043 | 0.00043 | 0.00021 | 1 | 2 |
| Luliconazole | 0.00063 | 0.00063 | 0.00031 | 1 | 2 |
| Miconazole | 0.36 | 0.089 | 0.71 | 4 | 0.5 |
| Neticonazole | 0.0094 | 0.0047 | 0.019 | 2 | 0.5 |
| Oxiconazole nitrate | 0.1 | 0.026 | 0.21 | 4 | 0.5 |
| Sulconazole | 0.26 | 0.033 | 0.52 | 8 | 0.5 |
| Imidazole pesticide |  |  |  |  |  |
| Dimetconazole | 0.2 | 0.0061 | 0.2 | 32 | 1 |
| Difenoconazole | 1.6 | 0.0061 | 1.6 | 256 | 1 |
| Tebuconazole | 1.6 | 0.024 | 0.78 | 64 | 2 |
| Hexaconazole | 1.6 | 0.012 | 0.78 | 128 | 2 |
| Myclobutanil | 3.1 | 0.098 | 3.1 | 32 | 1 |
| Oxpoconazole | 0.78 | 0.012 | 0.78 | 64 | 1 |
| Triflumizole | 3.1 | 0.024 | 3.1 | 128 | 1 |
| Propiconazole | 1.6 | 0.0061 | 1.6 | 128 | 1 |
| Pefurazoate | 0.2 | 0.1 | 0.1 | 2 | 2 |
| Imazalil | 0.13 | 0.0041 | 0.13 | 32 | 1 |
| Prochloraz | 0.26 | 0.13 | 0.032 | 2 | 8 |
| Others |  |  |  |  |  |
| Triforine | >100 | 25 | >100 | >4 | ND |
| Fenarimol | 1.6 | 0.049 | 1.6 | 32 | 1 |

### Synergistic Effects of Azole Combinations

Based on the different azole sensitivities of Δ*cyp51A* and Δ*cyp51B* strains, we explored the potential synergistic effects of azole combinations. We selected prochloraz, which demonstrated the highest fold changes in MIC values against Δ*cyp51B* strains, and fluconazole, sulconazole, and imazalil, which exhibited relatively high fold changes in MIC values, for the combination studies. In the WT strain, the combination resulted in fractional inhibitory concentration (FIC) indices of 0.32, 0.33, and 0.29 for the combination with fluconazole, sulconazole, and imazalil, respectively (Table 2), indicating synergistic effects. The Δ*ku80* strain demonstrated similar FIC indices (0.35, 0.28, and 0.31, respectively; Table 2), whereas the FIC indices for Δ*cyp51A* and Δ*cyp51B* strains were 1.0 and 0.62 for fluconazole, 0.81 and 0.78 for sulconazole, and 0.92 and 0.85 for imazalil, respectively (Table 2). These data suggest that dual azole combinations could serve as an effective therapeutic strategy against fungi expressing multiple Cyp51 isozymes.

**Table 2.** FIC index of prochloraz combined with imazalil, fluconazole, or sulconazole against *Trichophyton rubrum* WT, parent strain Δ*ku80*, Δ*cyp51A*, and Δ*cyp51B*. mean ± SD.

|  | FIC index |  |  |
| --- | --- | --- | --- |
|  | Fluconazole | Sulconazole | Imazalil |
| WT | 0.33 ± 0.14 | 0.29 ± 0.071 | 0.32 ± 0.12 |
|  | (n = 3) | (n = 3) | (n = 8) |
| $\Delta ku80$ | 0.28 ± 0.13 | 0.31 ± 0.13 | 0.35 ± 0.035 |
|  | (n = 2) | (n = 2) | (n = 4) |
| $\Delta cyp51A$ | 1.0 ± 0.0 | 0.81 ± 0.27 | 0.92 ± 0.51 |
|  | (n = 2) | (n = 2) | (n = 3) |
| $\Delta cyp51B$ | 0.62 ± 0.0 | 0.78 ± 0.31 | 0.85 ± 0.28 |
|  | (n = 2) | (n = 2) | (n = 3) |

To investigate whether the combination of azole antifungal drugs exerts a synergistic effect on other fungal species that possess Cyp51A and Cyp51B in their genome, we tested these combinations against *Aspergillus* sp., which also possesses two Cyp51 isozymes. The combinations of prochloraz with fluconazole, sulconazole, and imazalil were tested against the *Aspergillus welwitschiae* strains IFM57545 (relatively azole-susceptible strain) and IFM63877 (relatively azole-resistant strain). The FIC indices for the IFM57545 and IFM63877 strains were 0.41 and 0.38 for fluconazole, 0.44 and 0.25 for sulconazole, and 0.38 and 0.28 for imazalil, respectively (Table 3), suggesting the synergistic effects of these combinations. These data support the potential of this strategy against other pathogenic fungi.

**Table 3.**
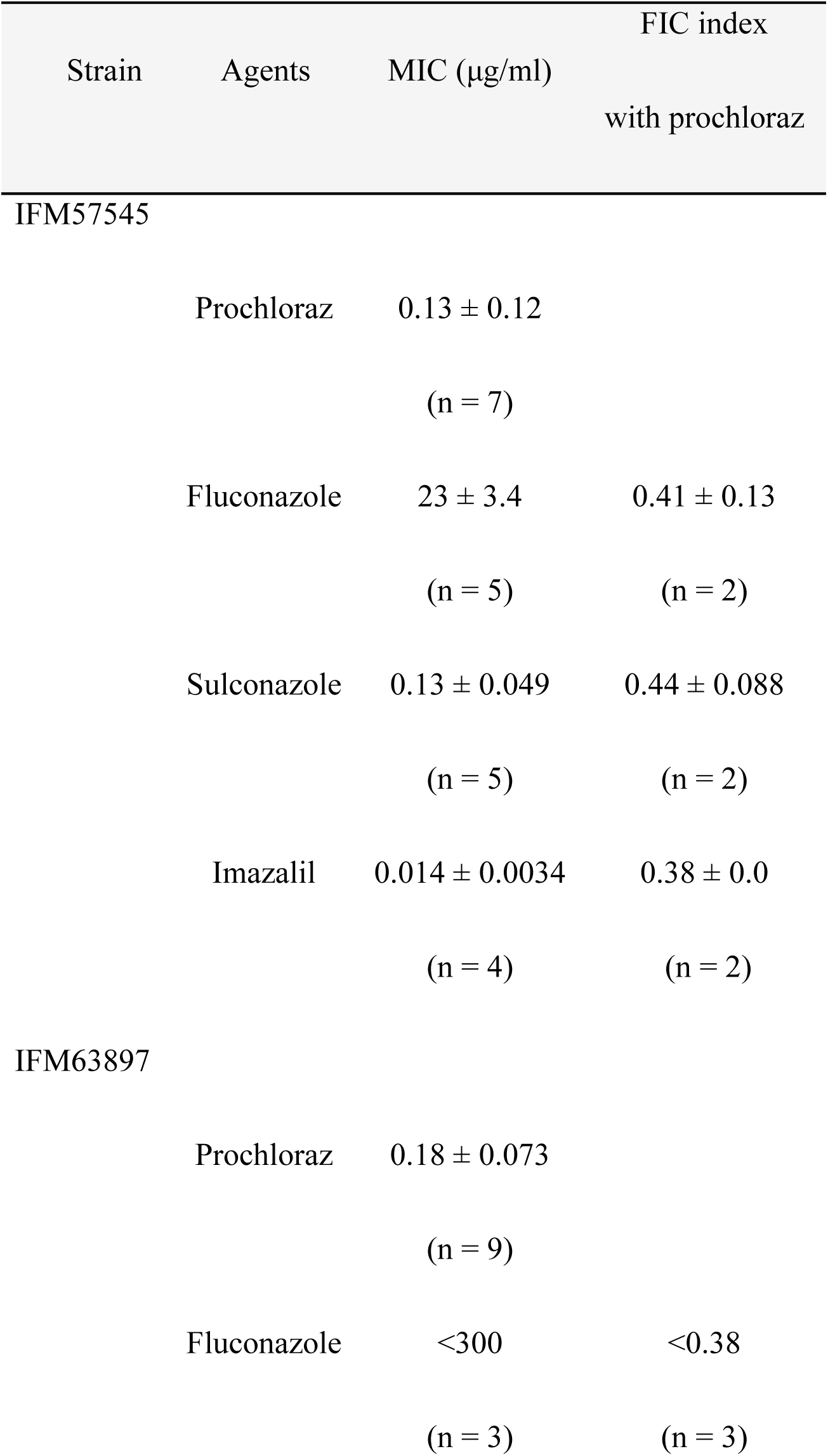
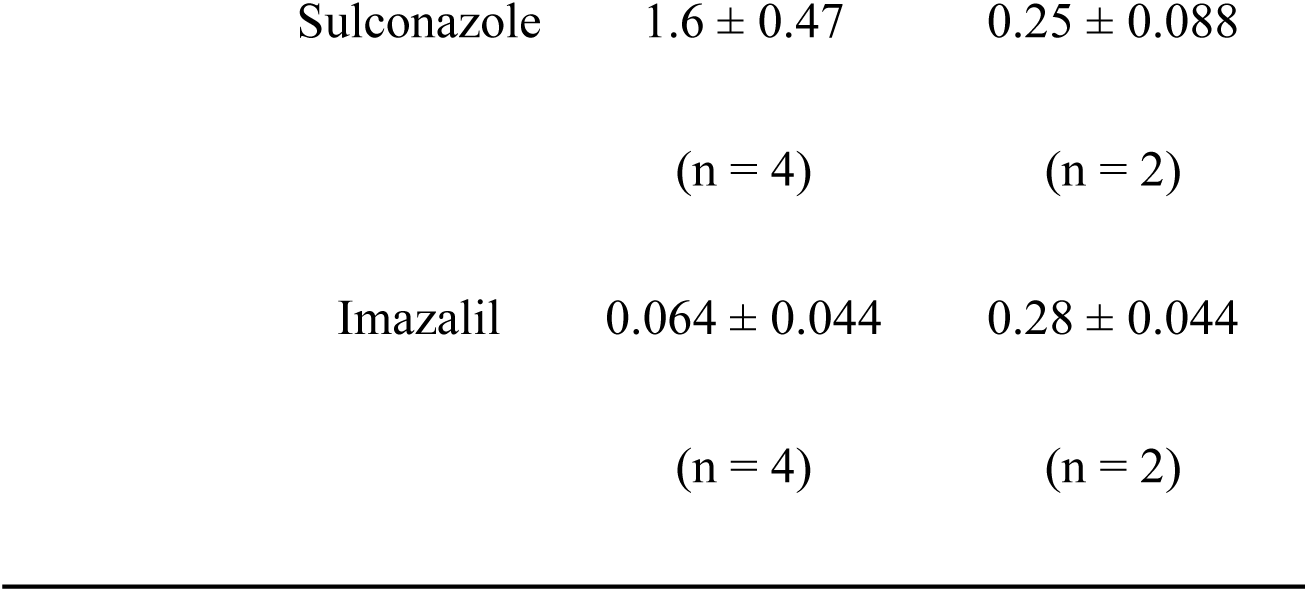
MIC values (μg/ml) ±SD and FIC index of prochloraz and imazalil combination in *A. welwitschiae* strains IFM57545 and IFM63897. mean ± SD.

| Strain | Agents | MIC ($\mu\text{g/ml}$) | FIC index<br>with prochloraz |
| --- | --- | --- | --- |
| IFM57545 |  |  |  |
| | Prochloraz | $0.13 \pm 0.12$<br>(n = 7) | |
| | Fluconazole | $23 \pm 3.4$<br>(n = 5) | $0.41 \pm 0.13$<br>(n = 2) |
| | Sulconazole | $0.13 \pm 0.049$<br>(n = 5) | $0.44 \pm 0.088$<br>(n = 2) |
| | Imazalil | $0.014 \pm 0.0034$<br>(n = 4) | $0.38 \pm 0.0$<br>(n = 2) |
| IFM63897 |  |  |  |
| | Prochloraz | $0.18 \pm 0.073$<br>(n = 9) | |
| | Fluconazole | $<300$<br>(n = 3) | $<0.38$<br>(n = 3) |

|  |  |  |
| --- | --- | --- |
| Sulconazole | $1.6 \pm 0.47$ | $0.25 \pm 0.088$ |
|  | (n = 4) | (n = 2) |
| Imazalil | $0.064 \pm 0.044$ | $0.28 \pm 0.044$ |
|  | (n = 4) | (n = 2) |

## DISCUSSION

This study demonstrated that the deletion of Cyp51 isozymes in *T. rubrum* significantly altered the organism’s susceptibility to azole antifungals. The differential MIC values observed in the Cyp51 isozyme deletion strains probably reflect the varying affinities of azole antifungals for these isozymes. This interpretation is supported by previous biochemical studies in *A. fumigatus*, where recombinant protein analysis revealed that Cyp51A exhibits 3.0– to 37-fold higher dissociation constants (K_d_) with azole antifungals compared with those exhibited by Cyp51B (16). Although these findings are similar to our observations regarding the predominant role of Cyp51A in azole tolerance, direct biochemical studies using recombinant *T. rubrum* Cyp51 isozymes will be required to confirm similar binding properties in this species.

Using isozyme-deficient mutants, we demonstrated that CYP51A and CYP51B exert distinct functions in *T. rubrum*. Cyp51B is essential for basal growth, as evidenced by the severe growth inhibition observed in the Δ*cyp51B* strain. In contrast, Cyp51A is not essential for basal growth but is induced under ergosterol depletion and promotes azole tolerance. This functional differentiation is remarkably different from observations in related fungi. For instance, the deletion of neither Cyp51A nor Cyp51B significantly affected mycelial growth in *A. fumigatus* (17). Similarly, *T. mentagrophytes*, a close relative of *T. rubrum*, exhibited minimal growth impairment after Cyp51B deletion (18). These contrasting findings suggest that the Cyp51 isozyme dependence of proliferation varies among species.

The synergistic effects observed with combinations of azoles possessing different isozyme selectivities have significant therapeutic implications. This finding not only suggests novel treatment strategies for dermatophytosis but may also be applicable to infections caused by other fungi with multiple Cyp51 isozymes, including *Aspergillus*, *Fusarium*, and potentially *Mucor* species, where drug resistance remains problematic (19). The differential response to azoles after isozyme deletion is not unique to *T. rubrum*; similar variations in azole susceptibility between isozymes have been reported in *Fusarium* species (20–23). Although a previous study reported synergistic effects between antifungal azoles (24), the underlying mechanisms remained unclear. Our study suggests that isozyme selectivity is a key factor driving these synergistic interactions. We observed that a considerable number of compounds demonstrated strong selectivity for the Cyp51B isozyme. Conversely, a limited number of compounds exhibited high specificity for the Cyp51A isozyme. In future studies, it will be necessary to identify compounds with high specificity for Cyp51A, which may in turn result in the discovery of combinations of compounds that exert even stronger synergistic effects.

This study opens novel avenues for the development of antifungal drugs. The development of screening systems focused on isozyme specificity could identify azole combinations with improved antifungal activity. Importantly, our results may revitalize interest in compounds previously dismissed during drug development due to weak overall antifungal activity, as these compounds may possess valuable isozyme-specific inhibitory properties that could be exploited in combination therapy approaches.

## MATERIALS AND METHODS

### Fungal and bacterial strains and culture conditions

*T. rubrum* CBS118892, obtained from Westerdijk Fungal Biodiversity Institute, was cultured on Sabouraud dextrose agar (SDA; 1% Bacto peptone, 4% glucose, 1.5% agar, pH unadjusted) at 28°C. *A. welwitschiae* IFM57545 and IFM63877 strains were obtained from NBRP (25). Conidia were prepared as described previously (26).

### Plasmid construction

To construct the *cyp51A*-targeting vector, pUC19Δ*cyp51A*, approximately 1.0 kb of the 5′-UTR fragments of the *cyp51A* open reading frame (ORF) was amplified from *T. rubrum* genomic DNA by PCR. The *neomycin phosphotransferase* gene cassette, which consists of *Escherichia coli* neomycin phosphotransferase (*nptII*), *A. nidulans trpC* promoter (*PtrpC*), and *A. fumigatus cgrA* terminator (*TcgrA*) with the 5′-UTR fragments of the *cyp51A* ORF, was amplified from pAg1-*cyp51A*-3′-UTR (15) using the primer pair *PtrpC*-F and *cyp51A*-3′-R-pUC19. The plasmid backbone of pUC19 was cleaved using *Kpn*I/PstI. These three fragments were joined using an In-Fusion HD Cloning Kit (TaKaRa Bio, Japan).

To construct the *cyp51B*-targeting vector, pUC19Δ*cyp51B*, approximately 1.4 and 1.1 kb of the 5′– and 3′-UTR fragments of the *cyp51B* ORF were amplified from *T. rubrum* genomic DNA. The plasmid backbone of pUC19 was cleaved using *Kpn*I/PstI. The *neomycin phosphotransferase* gene cassette, which consists of *nptII*, *PtrpC*, and *TcgrA*, was amplified from pAg1-ΔTr*cla4* (27). These fragments were joined using an In-Fusion HD Cloning Kit (TaKaRa Bio, Japan). The primers used in this study are listed in Table S1.

### Transformation of *T. rubrum*

*T. rubrum* was transformed using the polyethylene glycol (PEG) method as described previously (28, 29). The desired transformants and purified genomic DNA were evaluated by PCR. Total DNA was extracted using the Quick-DNA Fungal/Bacterial Miniprep Kit (Zymo Research, USA). Fungal cells were clashed using μT-01 (TAITEC, Japan) using 5-mm stainless beads.

### Antifungal susceptibility assay

Conidia (2 × 10^3^) were incubated with twofold serial dilutions of antifungal agents in 200 μl of MOPS-buffered RPMI (pH 7.0) at 28°C for 1 day (for *A. welwitschiae*) or 7 days (for *T. rubrum*), and MIC_100_ (concentration required to inhibit growth by 100%) was determined (30). Efinaconazole and luliconazole were purchased from BLD Pharmatech Ltd, China. Ravuconazole, isavuconazole, and triforine were purchased from Merck USA. Itraconazole, fluconazole, voriconazole, posaconazole, dimetconazole, ketoconazole, myclobutanil, difenoconazole, propiconazole, tebuconazole, hexaconazole, imazalil, and prochloraz were purchased from Tokyo Chemical Industry Co., Ltd, Japan. Oxpoconazole, clotrimazole, miconazole, triflunizole, pefurazoate, and fenarimol were purchased from FUJIFILM Wako Pure Chemical Corporation, Japan. Isoconazole and bifonazole were purchased from Thermo Fisher Scientific Inc.,USA. Sulconazole was purchased from Cayman Chemical Company, USA. Lanoconazole, neticonazole, and oxiconazole nitrate were purchased from MedChem Express, USA.

The FIC index was calculated using the following formula: FIC index = (MIC_Acom_/MIC_A_) + (MIC_Bcom_/MIC_B_), where MIC_Acom_ and MIC_Bcom_ are the MIC of drugs tested in combination, and MIC_A_ and MIC_B_ are the MIC of drugs tested alone. Synergy was defined as an FIC index of ≤0.5 (31). Each experiment was performed at least twice.

### Quantitative reverse transcription-PCR (qRT-PCR)

Total RNAs were purified using NucleoSpin RNA (Macherey-Nagel) and reverse-transcribed into cDNAs using ReverTra Ace (Toyobo) according to the manufacturers’ instructions. qRT-PCR was performed using TB Green Premix Ex Taq II (TaKaRa Bio, Japan) on a StepOne Real-time PCR system (Thermo Fisher Scientific, USA). The relative mRNA expression level was determined using the 2^-*ΔΔCt*^ method using *Chitin synthase I* (*chs1*) as an endogenous control to normalize the samples (29). The primers used in this study are listed in Table S1.

### Statistics

The mean values of three groups with one variable were compared using a one-way analysis of variance with Tukey’s post hoc test using Prism 9 (GraphPad, USA). Differences were considered significant at *P* < 0.05.

## • Acknowledgments

The authors thank H. Uga, N. Hori, K. Hasegawa, and Y. Nittono for their technical help.

## • conflicts of interest (COI)

The authors declare no conflicts of interest.

## • Funding

This work was partly supported by a DAIGAKUTOKUBETSU KENKYUHI Grant from Musashino University, the Takeda Science Foundation and a JSPS KAKENHI Grant.

## • Data availability

The data that support the findings of this study are available from the corresponding author upon reasonable request.

## Figure legend

**Table S1.**
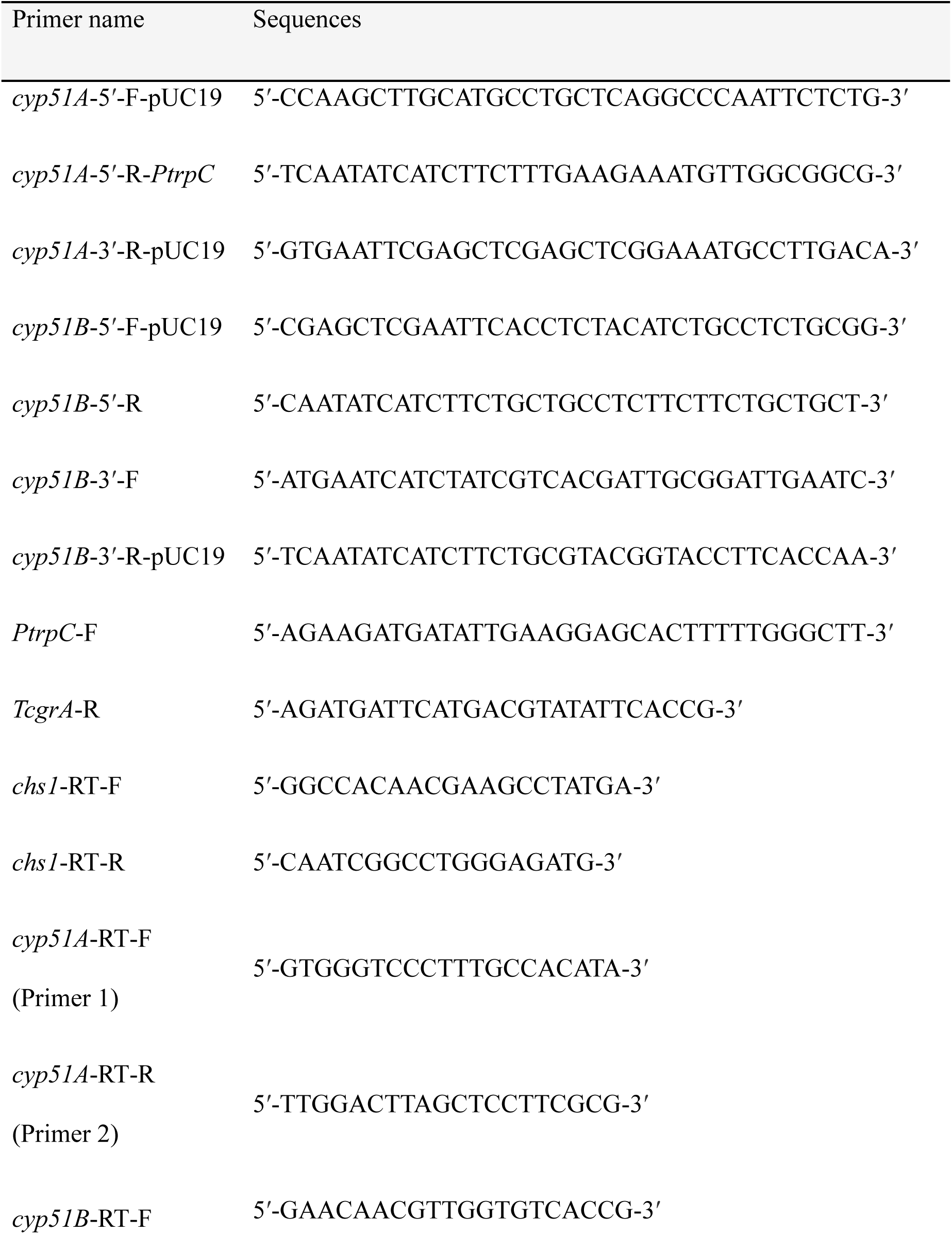
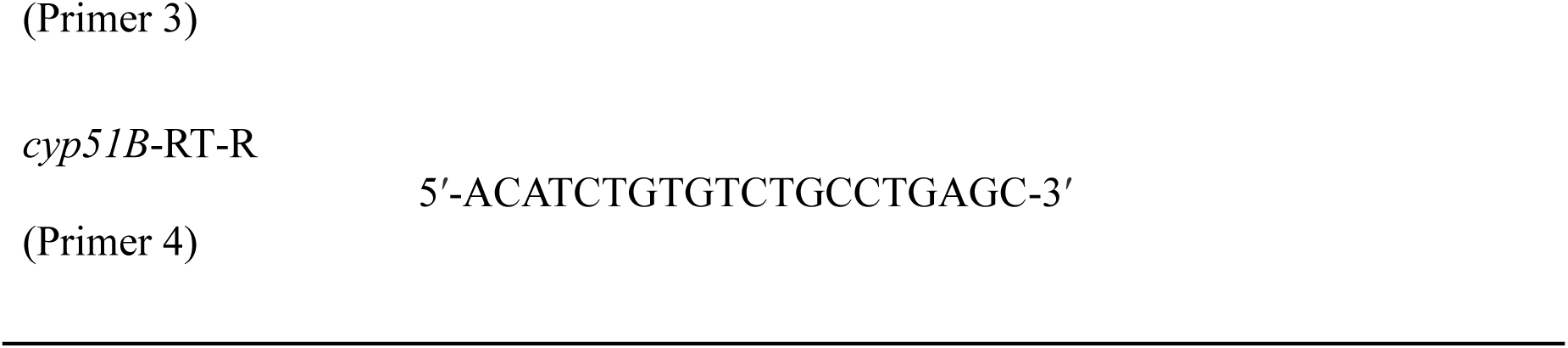
Primers used in this study.

